# A poxvirus decapping enzyme localizes to mitochondria to regulate RNA metabolism and translation, and promote viral replication

**DOI:** 10.1101/2021.10.22.465448

**Authors:** Shuai Cao, Joshua A Molina, Fernando Cantu, Candy Hernandez, Zhilong Yang

## Abstract

Decapping enzymes remove the 5’-cap of eukaryotic mRNA, leading to accelerated RNA decay. They are critical in regulating RNA homeostasis and play essential roles in many cellular and life processes. They are encoded in many organisms and viruses, including vaccinia virus, which was used as the vaccine to eradicate smallpox. Vaccinia virus encodes two decapping enzymes, D9 and D10, that are necessary for efficient viral replication and pathogenesis. However, the underlying molecular mechanism regulating vaccinia decapping enzymes’ function is still largely elusive. Here we demonstrated that vaccinia D10 localized almost exclusively to mitochondria that are highly mobile cellular organelles, providing an innovative mechanism to concentrate D10 locally and mobilize it to efficiently decap mRNAs. As mitochondria were barely present in “viral factories,” where viral transcripts are produced, suggesting that mitochondrial localization provides a spatial mechanism to preferentially decap cellular mRNAs over viral mRNAs. We identified three amino acids responsible for D10’s mitochondrial localization. Loss of mitochondrial localization significantly impaired viral replication, reduced D10’s ability to resolve RNA 5’-cap aggregation during infection, diminished D10’s gene expression shutoff and mRNA translation promotion abilities.

**Importance:** Decapping enzymes comprise many members from various organisms ranging from plants, animals, and viruses. The mechanisms regulating their functions vary and are still largely unknown. Our study provides the first mitochondria-localized decapping enzyme, D10, encoded by vaccinia virus that was used as the vaccine to eradicate smallpox. Loss of mitochondrial localization significantly impaired viral replication and D10’s gene expression shutoff and mRNA translation promotion ability. Mitochondrial localization is a spatial mechanism to concentrate D10 locally and mobilize it to efficiently and preferentially target cellular mRNAs for decapping and promote viral mRNA translation. Our results have broad impacts on understanding the functions and mechanisms of decapping enzymes.

## Introduction

The methyl guanosine cap (m^7^G) at the 5’-end of eukaryotic mRNA regulates many aspects of RNA processing and metabolism, such as splicing, transportation to the cytoplasm, protecting mRNA from 5’-3’ degradation by exonucleases, and recruiting cap-dependent translation initiation factors [1]. Decapping enzymes are proteins with many known members that regulate mRNA stability through removing the 5’-cap to render RNA sensitive to exonuclease-mediated 5’-3’ digestion. They are critical for regulating the homeostasis of cellular mRNAs levels and play crucial roles in numerous cellular and life processes [2]. In humans and various model systems, decapping enzymes are involved in cell migration, development, and cancers [3–7]. The active enzymatic activity of decapping enzymes lies in the Nudix motif with hydrolase activity, which hydrolyzes nucleoside diphosphate linked to other moieties [8]. The Nudix motifs are highly conserved and usually located in the center regions of the decapping enzymes [8]. Positive and negative regulatory domains are typically presented at the N- and C-termini of the proteins, which bind to either RNAs or other proteins to regulate the substrate specificities and potentials of decapping enzymes [9–12].

Dcp2 was the first discovered decapping enzyme from budding yeast *Saccharomyces cerevisiae* [13], followed by numerous homologs found in other organisms, including humans and plants [4, 14–17]. The human genome encodes multiple decapping enzymes [18]. Dcp2 carries out its catalytic activity in cytoplasmic structures called Processing bodies (P-bodies) [15]. P-bodies are cytoplasmic ribonucleoprotein (RNP) granules containing proteins involved in RNA degradation, including decapping enzymes, exonuclease Xrn1, and proteins involved in the RNA interference pathway beside translationally repressed mRNAs [19]. Local concentrating Dcp2 and other related proteins in P-bodies increases RNA degradation potential [20]. However, different decapping enzymes likely have distinct substrate specificities and modes of action. For example, Nudt16 is a nuclear decapping enzyme with a high affinity for U8 small nucleolar RNA [21]. Nudt12 is a cytoplasmic decapping enzyme that targets NAD+ capped RNA[22]. Today, while much progress has been made to understand decapping enzymes, how they achieve different functions and their molecular mechanisms of action remain largely elusive.

Interestingly, many viruses encode decapping enzymes, including poxviruses, Africa swine fever virus, and many other large nucleo-cytoplasmic DNA viruses [23–26]. Vaccinia virus (VACV), the vaccine used to eradicate the historically one of the most (if not the most) devastating infectious diseases, smallpox, encode two decapping enzymes, D9 and D10 [24, 25]. VACV is the prototypic member of the poxviruses, a large family of double-stranded DNA viruses currently causing many severe diseases in humans and economically and ecologically important animals [27, 28]. Poxviruses are also actively developed for treating cancers and as vaccine vectors [28]. D10 is present in all sequenced poxviruses [25]. They have low similarities to human Dcp2. D9 and D10’s decapping activities were demonstrated *in vitro* [24, 25]. They negatively regulate viral and cellular gene expression in VACV infected cells by accelerating mRNA turnover and are thought to be critical to controlling the VACV cascade gene expression program to ensure sharp transitions [29–33]. However, it is unclear if these viral decapping enzymes employ mechanisms to preferentially target cellular mRNAs in VACV-infected cells. VACV infection produces excessive RNAs, and some of them can form dsRNA to stimulate receptor 2′5′-oligoadenylate synthetase 2 (OAS)-RNase L pathway and PKR activation, which lead to RNA decay and mRNA translation repression, respectively. D9 and D10 are among the essential viral factors to resolve excessive double-stranded RNA (dsRNA) produced in VACV infection to evade these antiviral immunities [34]. Our recent data identified another function of D9 and D10, which are required for efficient VACV mRNA translation during infection [35]. Strikingly, D10 alone promotes viral mRNA translation in uninfected cells to ensure high levels of viral protein production [35]. Promoting mRNA translation by D10 is unusual as decapping enzymes are thought to negatively regulate RNA translation by competing with cap-binding translation initiation factors [19, 36–39]. However, how D10 promotes mRNA translation is still largely unknown.

Here we demonstrated that D10 located almost exclusively to mitochondria, the first discovered among known decapping enzymes. We further identified the amino acids at the N-terminus of D10 that are critical for D10’s mitochondrial localization. The mitochondrial localization of D10 is required for efficient viral replication, and D10’s ability to regulate mRNA metabolism, translation promotion. The results indicate that mitochondria riding provides D10 a mechanism to concentrate proteins locally with remarkable mobility to preferentially decap cellular mRNAs during VACV infection.

## Results

### VACV D10 localizes to mitochondria

We examined D10’s subcellular localization using a recombinant VACV vD10-3xFlag, in which D10 was tagged with a 3xFlag epitope at the C-terminus. We used primary human foreskin fibroblasts (HFFs) and HeLa cells and found that D10 almost exclusively localizes to mitochondria (**Fig 1AB, Fig S1**). A549DKO human lung carcinoma cell, in which the PKR and RNase L genes were knocked out via CRISPR/Cas9, is very useful in studying VACV decapping enzyme functions. It excludes the PKR and RNase L activation-related RNA degradation and translation repression during decapping enzymes-inactivated VACV infection [34]. Again, D10 localized to mitochondria during infection as it colocalized with MitoTracker and Tom20, a well-known mitochondrial protein [40] (**Fig 1C**, **Fig. S2**). Another notable observation is that mitochondria barely reside in the viral factories (the cytoplasmic sites of viral replication with intensive DNA staining of viral DNA) (**Fig 1 A-C**). Using a plasmid expressing D10 with a C-terminal 3xFlag, we observed D10 localized to mitochondria in uninfected A549DKO and HeLa cells (**Fig 1D**). Together, our results demonstrate that VACV D10 localizes to mitochondria either during viral infection or in uninfected cells.

**Fig 1.**
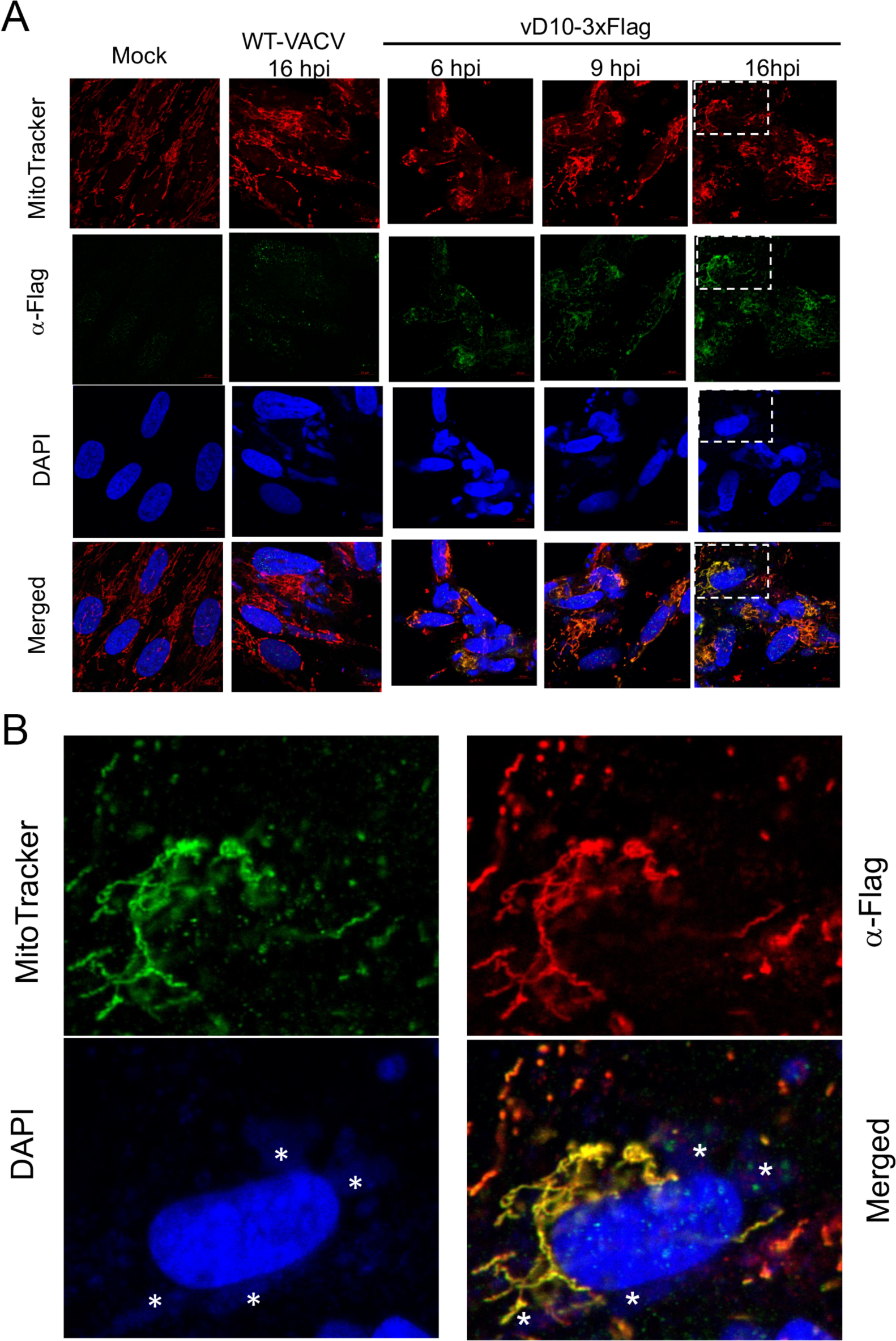
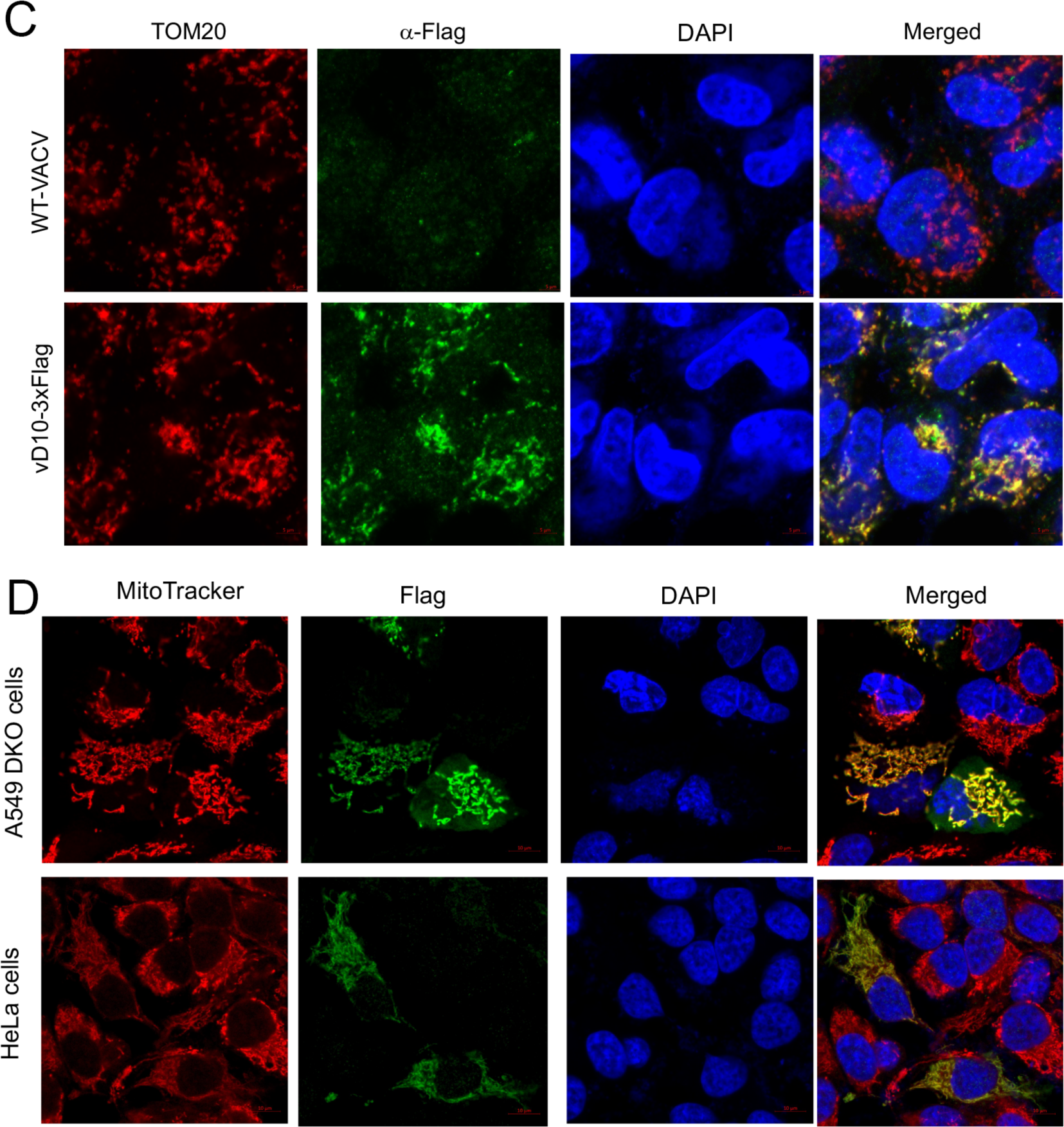
VACV D10 localizes to mitochondria. **(A)** D10 localizes to mitochondria in HFFs during VACV infection. HFFs were infected with vD10-3xFlag, or WT-VACV (MOI=3), or mock-infected. Confocal microscopy was used to visualize D10 (*α*-Flag antibody, green), mitochondria (MitoTracker, red), and DNA (DAPI, blue) at 6, 9, and 16 hpi (hours post-infection). **(B)** Zoomed in the indicated areas in A. The asterisks (*) indicate viral factories. **(C)** D10 localizes to mitochondria in A549DKO cells during VACV infection. A549DKO cells were infected with vD10-3xFlag or WT-VACV (MOI=3). Confocal microscopy was employed to visualize D10 (anti-Flag antibodies, green), mitochondria (*α*-Tom20 antibody, red), DNA (DAPI, blue) at 16 hpi. **(D)** D10 localizes to mitochondria in uninfected cells, A549 DKO or HeLa cells were transfected with plasmid encoding codon-optimized D10 with a C-terminal 3xFlag tag. Confocal microscopy was used to visualize D10 (*α*-Flag antibody, green), mitochondria (MitoTracker, red), and DNA (DAPI, blue) at 24 hours post-transfection.

### Identification of N-terminal hydrophobic amino acids required for D10 mitochondrial localization

We first generated and tested plasmids expressing a D10 C-terminal (D10-ΔC57) and a D10 N-terminal (D10-ΔN50) truncation mutants, respectively (**Fig 2A**). While D10-ΔC57 remained to localize to mitochondria, the D10-ΔN50 lost its mitochondrial localization ability (**Fig 2B**), suggesting that the N-terminal amino acids are required for D10 mitochondrial localization. Further truncations of D10 indicated that the N-terminal amino acids from 9 to 13 are needed for D10 localization to mitochondria because deletion of the first eight N-terminal amino acids did not fully block D10 mitochondria localization. In contrast, deletion of the first N-terminal 13 amino acids rendered D10 loss its mitochondrial localization (**Fig 2A, Fig S3**). We further tested two additional D10 mutants: D10-Δ9-13, in which the three amino acids from 9-13 (ISQII) were deleted, and D10-I9/12/13T, in which the hydrophobic Isoleucine (I) at 9, 12, 13 were changed to Threonine (T, neutral) (**Fig. 2A**). The rationale of the latter is that those hydrophobic residues that can form helix may interact with mitochondrial proteins or target mitochondrial membrane to dock D10 on mitochondria. The deletion mutant (D10-Δ9-13) largely, while the point mutation mutant (D10-I9/12/13T) entirely rendered D10 to lose its mitochondrial localization in uninfected cells (**Fig 2B**).

**Fig. 2.**
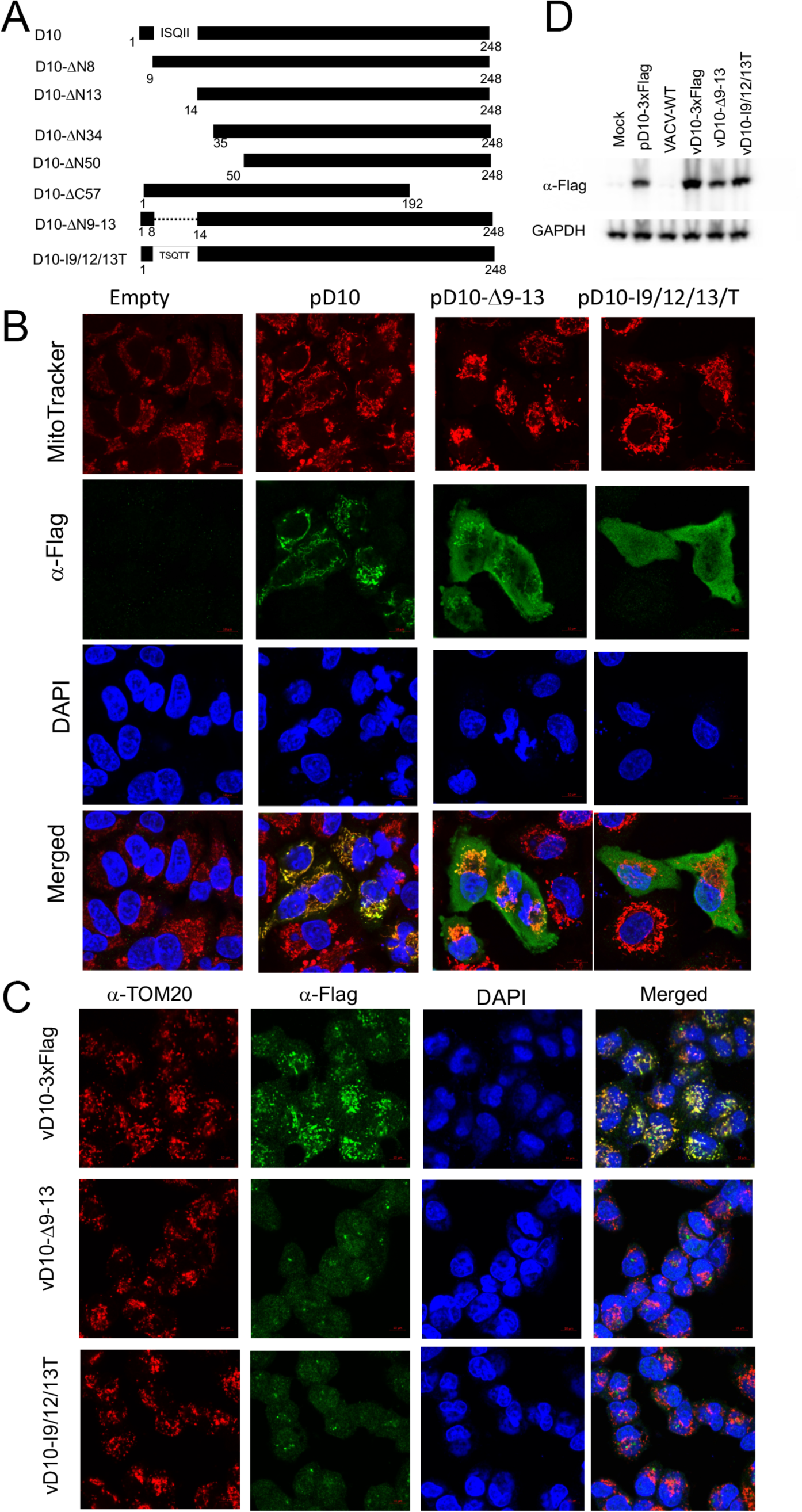
Three Isoleucines located at the N-terminal hydrophobic region of D10 are required for its mitochondrial localization. **(A)** Schematic of D10 mutants used in this study. **(B)** The hydrophobic amino acids Isoleucines located at the N-terminus of D10 are critical for D10 localization to mitochondria. A549 DKO cells were transfected with a plasmid expressing indicated codon-optimized D10 truncation mutants with a C-terminal 3xFlag. Confocal microscopy was employed to visualize D10 or its mutants using *α−*Flag antibody (green), mitochondria (MitoTracker, red), and DNA (DAPI, blue) at 24 h post-transfection. **(C)** D10 with amino acids 9-13 deletion or mutation expressed from recombinant VACV does not localize to mitochondria during infection, A549DKO cells were infected with indicated recombinant VACVs (MOI=3) encoding D10 mutants with a C-terminal 3xFlag tag. Confocal microscopy was used to visualize D10 (*α*-Flag antibody, green), mitochondria (*α*-Tom20, red), and DNA (DAPI, blue) at 16 hpi. **(D)** The levels of D10 or its mutants expressed from recombinant VACVs are expressed at comparable levels. A549DKO cells were infected with indicated viruses (MOI=3) or mock-infected. Western blotting analysis was employed to examine 3xFlag-tagged D10 expression using *α*-Flag antibody.

We then constructed two recombinant VACVs: vD10-I9/12/13T and vD10-Δ9-13, in which the D10 amino acids from 9 to 13 (ISQII) were mutated to TSQTT or deleted, respectively, yet both contained 3xFlag tag at the C-terminal. Interestingly, in both cases, the mutated D10 diffused in the infected A549 DKO or HeLa cells but did not localize to mitochondria, using Tom20 or Mitotracker to stain the mitochondria (**Fig 2C, Fig S4, Fig S5**). In addition, Western blotting analysis showed comparable protein levels of D10 and its mutants produced from the recombinant viruses (**Fig 2D**). These results corroborate that the N-terminal amino acids ISQII are required for D10 localization to mitochondria.

### Loss of D10 mitochondrial localization impairs VACV replication

Next, we examined the impact of D10 mitochondrial localization on VACV replication by comparing the replication of vD10-Δ9-13 and vD10-I9/12/13T to vD10-3xFlag, a control VACV encoding wild-type (WT) D10 with a 3xFlag tag at its C-terminus. vΔD10 is a recombinant VACV with D10 knocked out, and vD10mu is a recombinant VACV with D10’s decapping enzyme inactivated by mutating its Nudix motif [31], were included in the experiments. We used both A549 control and A549DKO cells as Liu et al. had shown that A549DKO cells could better support decapping enzyme inactivated VACV replication [34]. The A549 control cells were generated in parallel with A549DKO cells but with no PKR and RNase L knocked out [34]. All the recombinant viruses with mutated or deleted D10 replicated at a lower rate for a multiplicity of infection (MOI) of 3 and 0.001. The vD10-I9/12/13T more closely mimicked vΔD10 with more severe effects (∼3-fold higher decrease in both A549 control and A549DKO cells: 6-vs. 2.5-fold at MOI of 3 and up to 15-vs. 5-fold at MOI of 0.001 at 4 hpi) on viral yields and replication kinetics than that of vD10mu and vD10-Δ9-13 (**Fig 3A-D**). In addition, the reductions of VACV replication for all the tested mutant viruses were more prominent in A549 control cells than in A549DKO cells (**Fig 3A-D**). Interestingly, in BHK-21 cells, the decrease of D10 mutant viruses was similar to that in A549DKO cells with only moderate effects (**Fig S6**).

**Fig 3.**
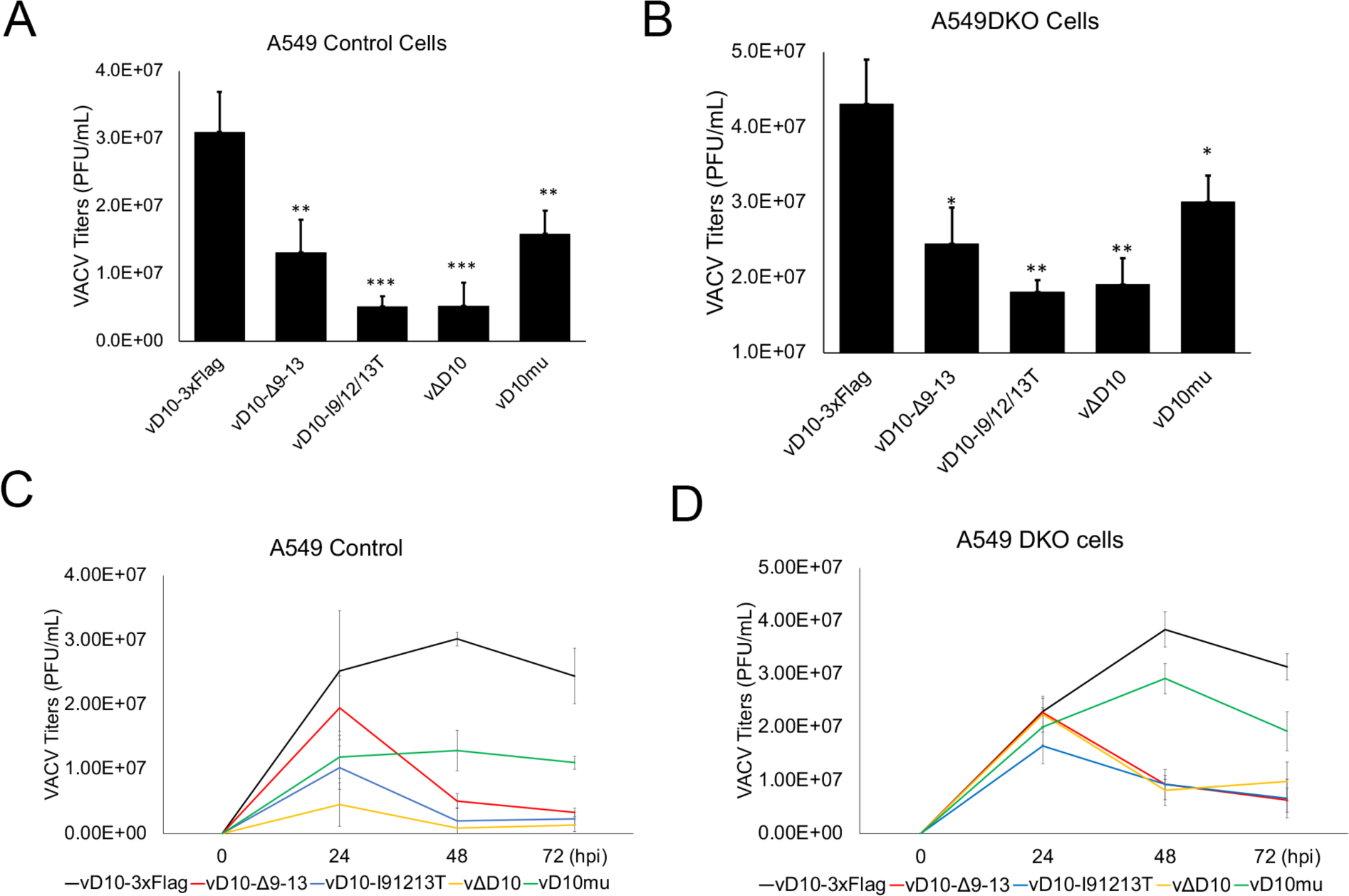
Loss of D10 mitochondrial localization impairs VACV replication in the presence of D9. **(AB)** A549 control **(A)** or A549DKO cells **(B)** were infected with indicated viruses at an MOI of 3. **(CD)** A549 control **(C** or A549DKO cells **(D)** were infected with indicated viruses at an MOI of 0.001. Viral titers were determined using a plaque assay at indicated times post-infection. All the viruses used encode D9. Error bars represent the standard deviation of at least three replicates. ns, P > 0.05; **, P ≤ 0.01; ***, P ≤ 0.001. Significance was compared to vD10-3xFlag.

VACV-encoded two decapping enzymes, D9 and D10, have overlapping functions [24, 25]. We rationalized that the loss of D10 mitochondrial localization has a more prominent effect on VACV replication in the absence of D9 expression. We generated a recombinant VACV vΔD9-D10-I9/12/13T, in which the D9 was knocked out and compared its replication with vΔD9 (D9 knocked out, wild type D10). We included vD10-3xFlag and vD9muD10mu (in which the decapping activities of both D9 and D10 are deactivated [32]) in this experiment. Consistent with a previous report [31], the replication of vΔD9 was not or only slightly affected in A549 control and A549DKO cells at MOI of 3 and 0.001, while vD9muD10mu barely replicated in A549 control cells but could replicate at some levels in A549DKO cells (**Fig 4A-D**). Notably, compared to vΔD9, vΔD9-D10-I9/12/13T showed an 83-fold and 11-fold reduction of viral yield at MOI of 3 in A549 control and A549DKO cells, respectively (Fig 4AB). At MOI of 0.001, vΔD9-D10-I9/12/13T replication also showed an 18-fold and 11-fold reduction of viral yields at its replication peaks in A549 control and A549DKO cells, respectively (Fig 4CD). A comparison of the plaque sizes indicated that vΔD9-D10-I9/12/13T and vΔD9-D10Δ9-13 have significantly smaller plaques than vD10-3xFlag (Fig 4EF). The plaque sizes of vΔD10 and vD10-I9/12/13T were also smaller than vD10-3xFlag (Fig 4EF). Overall, we conclude that D10 mitochondrial localization is required for efficient VACV replication in both PKR-and RNase L-dependent and independent manners.

**Fig. 4.**
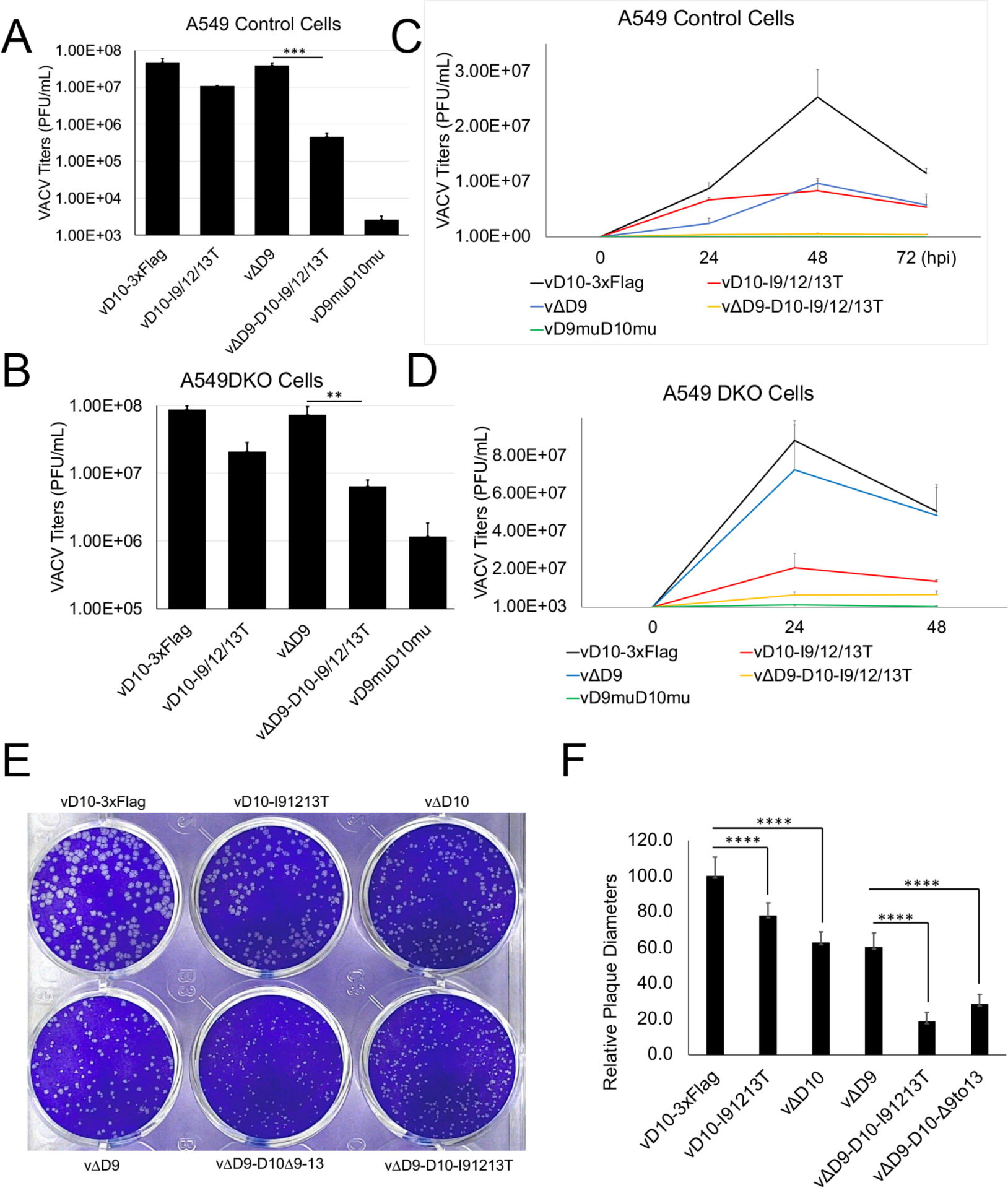
Loss of D10 mitochondrial localization significantly impairs VACV replication in the absence of D9 expression. **(A-D)** A549 control **(AC)** or A549DKO cells **(BD)** were infected with indicated viruses at an MOI of 3 **(AB)** or 0.001**(CD).** D9 was knocked out in vΔD9 and vΔD9-D10-I9/12/13T. Viral titers were determined using plaque assay at indicated times post-infection. Error bars represent the standard deviation of at least three replicates. **(E)** Loss of D10 mitochondrial localization reduced VACV plaque sizes. BS-C-1 cells were infected with indicated viruses. Plaques were visualized by a plaque assay. **(F)** Diameters from 25 plaques were measured using Image J and plotted. The diameters of vD10-3xFlag plaques were normalized to 100. **, P ≤ 0.01; ***, P ≤ 0.001; ****, P ≤ 0.0001.

### Loss of D10 mitochondrial localization reduces viral protein production during VACV infection

Our results (**Figs 3&4**) demonstrated that the effect of D10 mitochondrial localization on VACV replication could be more readily observed in the absence of D9. We then compared viral protein expression levels of vΔD9, vΔD9-D10-I9/12/13T, and vD9muD10mu during infection. While more or similar levels of viral early (E3) and intermediate (D13) proteins from vΔD9-D10-I9/12/13T and vD9muD10mu to that from vΔD9 infection were detected before 8 hpi, we observed substantially less intermediate (D13) and late (A7) viral protein levels at 16 and 24 hpi from vΔD9-D10-I9/12/13T and vD9muD10mu infection (**Fig 5AE**). Similarly, we observed substantially less total viral protein production from vΔD9-D10-I9/12/13T and vD9muD10mu infection than vΔD9 infection (**Fig 5AE**). There are two other notable observations: (1) the reduction of viral protein production during late infection was more prominent, (2) the extent of protein production reduction was less in ΔD9-D10-I9/12/13T than in vD9muD10mu infection (**Fig 5AE**).

**Fig. 5.**
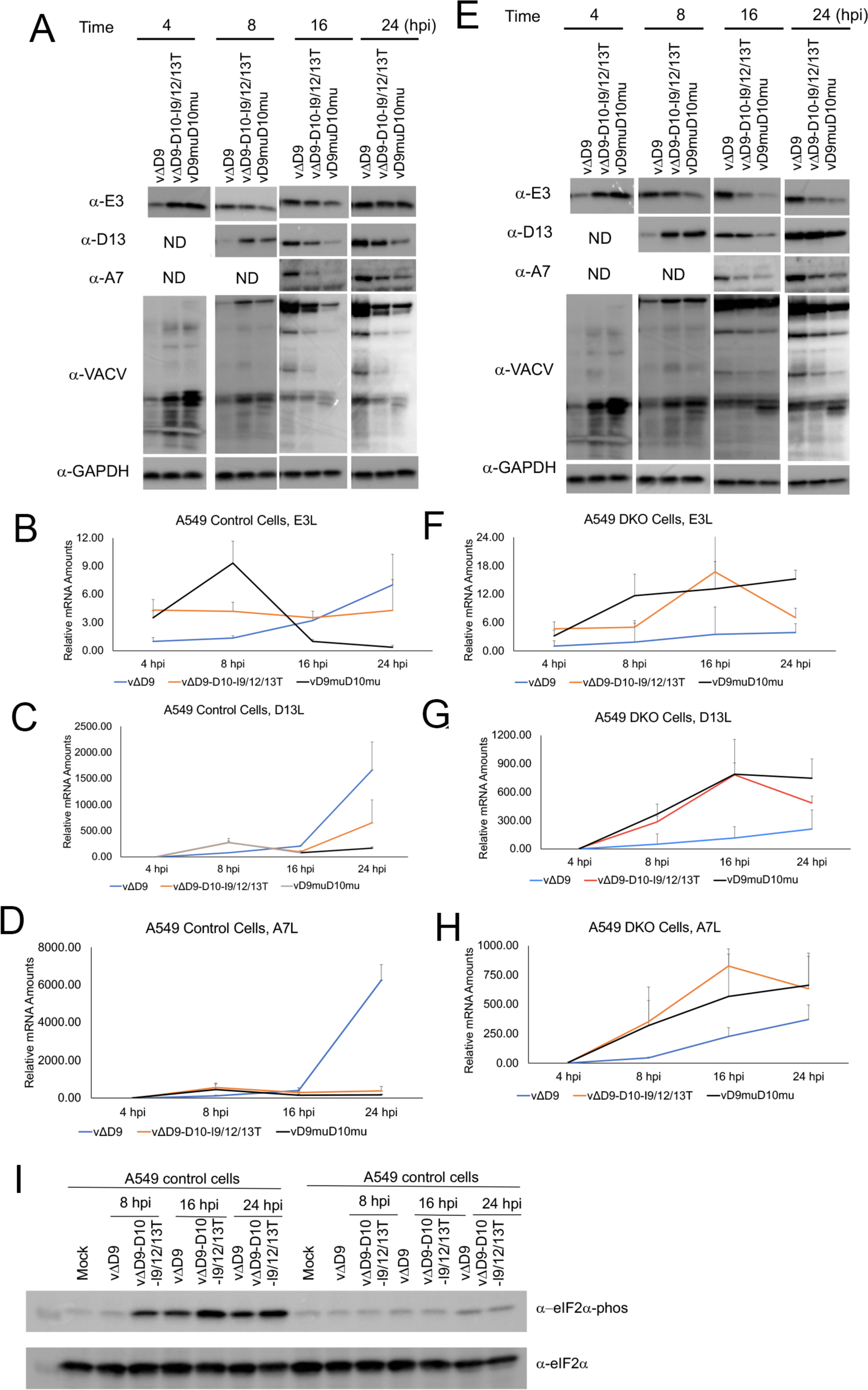
Loss of D10 mitochondrial localization reduces viral protein production during VACV infection. **(AE)** A549 control **(A)** or A549DKO cells **(E)** were infected with indicated viruses at an MOI of 3. Viral proteins were detected using indicated antibodies at indicated times post-infection. GAPDH was used as a loading control. E3, D13, and A7 are viral early, intermediate, and late proteins, respectively. **(BCD)** A549 control cells were infected with indicated viruses at an MOI of 3. Relative levels (to cellular 18S) of viral mRNAs were quantified using qRT-PCR. **(B)** E3L (early), **(C)** D13L (intermediate), **(D)** A7L (late). The mRNA levels were normalized to the level at 4 hpi in vΔD9-infected cells for each mRNA. **(FGH)** A549 control cells were infected with indicated viruses at an MOI of 3. Relative levels (to cellular 18S) of viral mRNAs were quantified using qRT-PCR. **(F)** E3L (early), **(G)** D13L (intermediate), **(H)** A7L (late). The mRNA levels were normalized to the level at 4 hpi in vΔD9-infected cells for each mRNA. Error bars represent the standard deviation of at least three replicates. **(I)** Western blotting analysis of eIF2*α* phosphorylation in A549 control and A549DKO cells infected with indicated viruses at indicated times post-infection.

To determine if the protein synthesis reduction was correlated to viral mRNA levels, we carried out quantitative real-time PCR (qRT-PCR) to measure E3L, D13L, and A7L mRNAs at 4, 8, 16, and 24 hpi. In A549 control cells, while the E3L, D13L, and A7L mRNA levels continued to increase over the course of VACV infection, a notable increase of these mRNAs was not observed in vΔD9-D10-I9/12/13T and vD9muD10mu infection. Notably, at 24 hpi, D13L and A7L mRNA levels in vΔD9-D10-I9/12/13T and vD9muD10mu infection were significantly lower than in vΔD9 infection (**Fig 5B-D**), likely due to the activation of RNase L RNA degradation pathway [34]. Interestingly, we observed generally higher E3L, D13L, and A7L mRNA levels in vΔD9-D10-I9/12/13T and vD9muD10mu infection than vΔD9 in A549DKO cells from 8 hpi (**Fig 5F-H**). It has been shown that inactivation of D9 and D10 decapping activities stimulates PKR activation, followed by eIF2*α* phosphorylation [32], leading to translation repression. We compared eIF2*α* phosphorylation in vΔD9 and vΔD9-D10-I9/12/13T infected cells and observed higher eIF2*α* phosphorylation in vΔD9-D10-I9/12/13T-infected A549 control cells but not A549DKO cells (**Fig 5I**), which likely contributed to the more severe protein production defect in A549 control cells (**Fig 5AB**). Since protein levels from vΔD9-D10-I9/12/13T infection were lower than that from vΔD9 infection, and PKR is not present in A549DKO cells (**Fig. 5E**), it suggests an additional translational disadvantage due to the loss of mitochondrial localization independent of PKR activation-induced translation suppression.

Together, these results demonstrate that loss of mitochondrial localization substantially decreases viral protein production in both A549 control and DKO cells. However, loss of mitochondrial localization leads to higher levels of viral mRNAs were observed in A549DKO cells but not in A549 control cells, again suggesting PKR- and RNase L-dependent and independent mechanisms.

### Loss of mitochondrial localization reduces D10’s gene expression shutoff ability

Both D9 and D10’s decapping activities are inactivated in vD9muD10mu [32]. In A549 control cells, the RNase L RNA degradation pathway was activated during VACV infection that explains lower viral mRNA levels in vD9muD10mu infection than in vΔD9 infection. However, in A549DKO cells, the knockout of RNase L inactivates the pathway, which leads to viral mRNA accumulation in vD9muD10mu infection [32, 34]. Our results in **Fig 5** indicate that vΔD9-D10-I9/12/13T closely mimics D9muD10mu for the effects on VACV mRNA levels, suggesting that the loss of mitochondrial localization may impair its ability to induce mRNA turnover. Because decapping enzymes remove 5’-m^7^G cap from mRNA, we employed an immunofluorescence assay to visualize mRNA 5’-caps in cells using *α*-cap antibodies. We used A549DKO cells as it supports the replication of VACV with inactivated decapping enzymes. In mock- and wild-type VACV-infected A549DKO cells, the caps distributed in the cells mostly evenly with no aggregation (**Fig 6A**). Strikingly, heavy aggregation of the 5’-caps was observed in almost all cells infected with vD9muD10mu. The aggregation was located primarily between the nuclei and viral factories (VACV DNA replication site in the cytoplasm with intensive DAPI staining) (**Fig 6A**). The aggregation of the cap indicates the inability of vD9muD10mu to remove RNA 5’-cap during infection. We then investigated the impacts of mitochondrial localization loss on m^7^G-cap aggregation in the presence or absence of D9 expression during VACV infection (**Fig. 6BC**). In the presence of D9, loss of D10 mitochondrial localization or D10 deletion slightly increased the number of cells with 5-cap aggregation, with 4% and 2.5% of cells with aggregation (**Fig 6B**). Notably, in the absence of D9 expression, the loss of D10 mitochondrial localization substantially increased the cap aggregation during infection, with 26% and 20% of cells for vΔD9-I9/12/13T and vΔD9-ΔD109-13, respectively (**Fig 6C**).

**Fig. 6.**
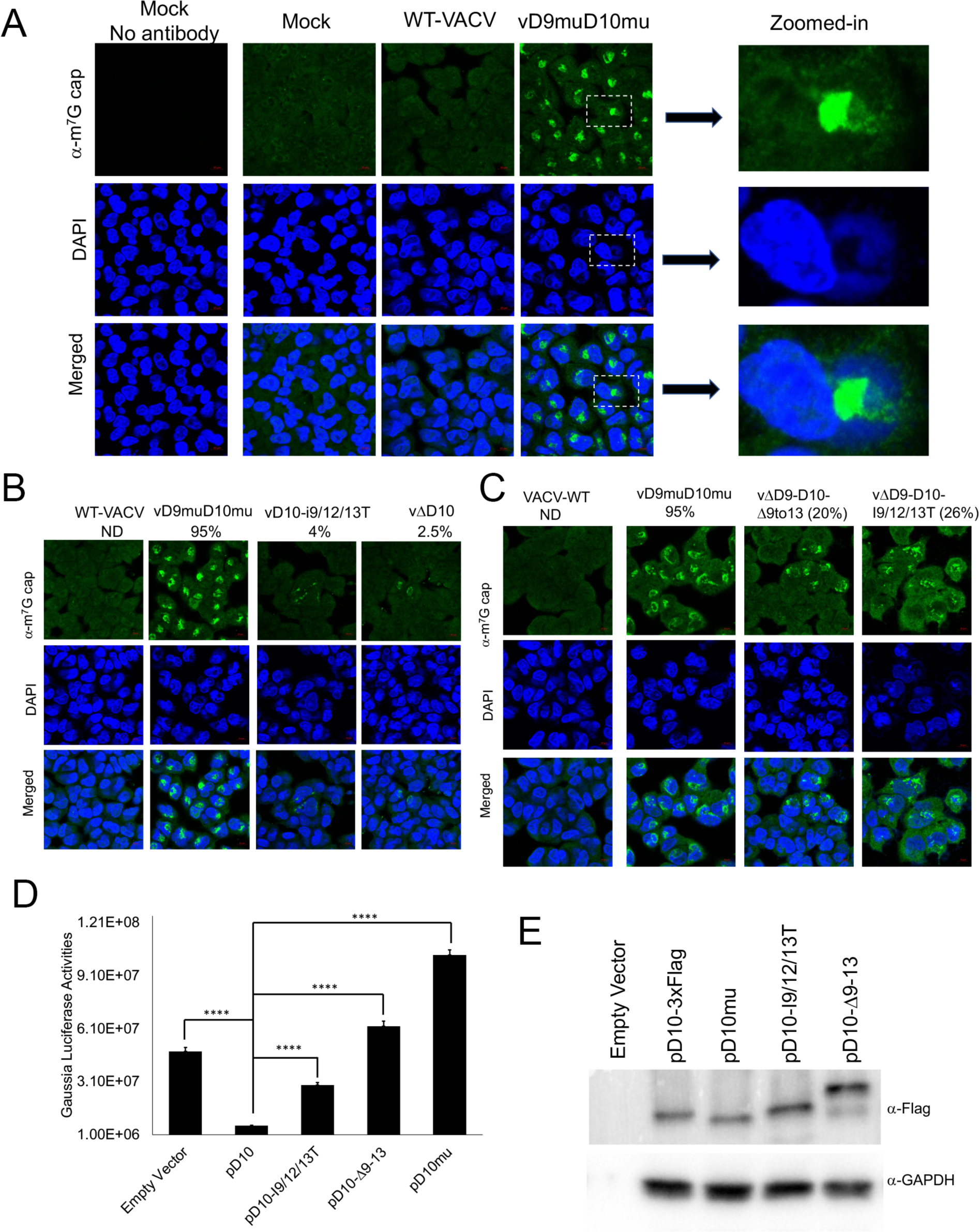
Loss of mitochondrial localization reduces D10’s gene expression shutoff ability. **(A)** Inactivation of D9 and D10’s decapping activities leads to aggregation of m^7^G cap structures in VACV-infected cells. A549DKO cells were infected with WT, vD9muD10mu (MOI=3), or mock-infected. Confocal microscopy was used to visualize m^7^G cap (*α*-cap antibody, green) and DNA (DAPI, blue) at 16 hpi. Three zoomed-in areas were shown on the right. The asterisks (*) indicate viral factories. **(BC)** Loss of mitochondrial localization leads to aggregation of the caps in VACV infected cells. A549 DKO cells were infected with indicated viruses at an MOI of 3. Confocal microscopy was used to visualize m^7^G cap (*α*-cap antibody, green) and DNA (DAPI, blue) at 16 hpi. The numbers indicate the percentages of cells with cap aggregation. **(D)** Loss of D10 mitochondrial localization reduces its ability to shut off gene expression. Gaussia luciferase reporter gene under a cellular EF-1a promoter was co-transfected with the indicated codon-optimized D10 or D10 mutants. Gaussia luciferase activities were measured 24 h post-transfection. **(E)** Western blotting analysis of D10 and D10 mutant protein levels. Error bars represent the standard deviation of at least three replicates. ****, P ≤ 0.0001.

These results (**Figs 5** **& 6B**) suggest that mitochondrial localization is required for efficient decapping in cells, which leads to RNA degradation and gene expression shutoff. We employed a virus-free approach to test if loss of mitochondrial localization impairs D10’s ability to induce gene expression shutoff in cells without interference from other viral factors. We co-transfected plasmids encoding codon-optimized D10 or its mutants with a Gaussia luciferase reporter plasmid under a cellular EF-1a promoter. As expected, WT D10 potently decreased Gaussia luciferase activity by 7.8-fold (**Fig 6D**). Very interestingly, co-transfection of a plasmid expressing D10-I9/12/13T could only reduce Gaussian luciferase expression by 1.7-fold (**Fig 6D**). D10Δ9-13 and D10mu (with Nudix domain mutation) lost their ability to suppress Gaussia luciferase expression (**Fig 6D**). The protein expression levels of D10 and its mutants from plasmids were comparable, although D10Δ9-13 showed a slower migration when expressed from a plasmid (**Fig 6E**). Taken together, our results indicate that loss of mitochondrial localization reduces D10’s ability to remove mRNA 5’-caps and shut off gene expression.

### Loss of mitochondrial localization impairs D10’s mRNA translation enhancement ability

D10 could promote mRNA translation, especially for mRNAs with a 5’-poly(A) leader, a feature of all poxvirus mRNAs expressed after viral DNA replication [35]. The enhancement is more notable for RNA without a 5’-m^7^G cap and could be revealed in the absence of VACV infection [35]. We employed an RNA-based luciferase reporter described previously [41, 42]. We used 293T cells in these experiments as we found this cell line had the highest transfection efficiency in the cells we tested. *In vitro* transcribed firefly luciferase (FLuc) RNA with a 5’-poly(A) leader and m^7^G-cap and renila luciferase (RLuc) RNA with a 5’-UTR containing Kozak sequence and m^7^G-capped were con-transfected in cells with expression of wild-type D10 or its mutants. Notably, all those containing mitochondrial localization sequence promoted 5’-poly(A) leader-mediated translation, while those without mitochondrial localization sequence significantly reduced the translation enhancement (**Fig. 7ABC).** ApppG-capped RNA translation only occurs in a cap-independent manner [43, 44]. The same trends were observed when ApppG-capped, 5’-poly(A) leader Fluc mRNA was used, although the translation enhancement was much higher than m^7^G-capped RNA (**Fig 7DEF**). Together, our results show that the mitochondrial localization is required for D10 to stimulate 5’-poly(A) leader mRNA translation, including cap-independent translation enhancement.

**Fig. 7.**
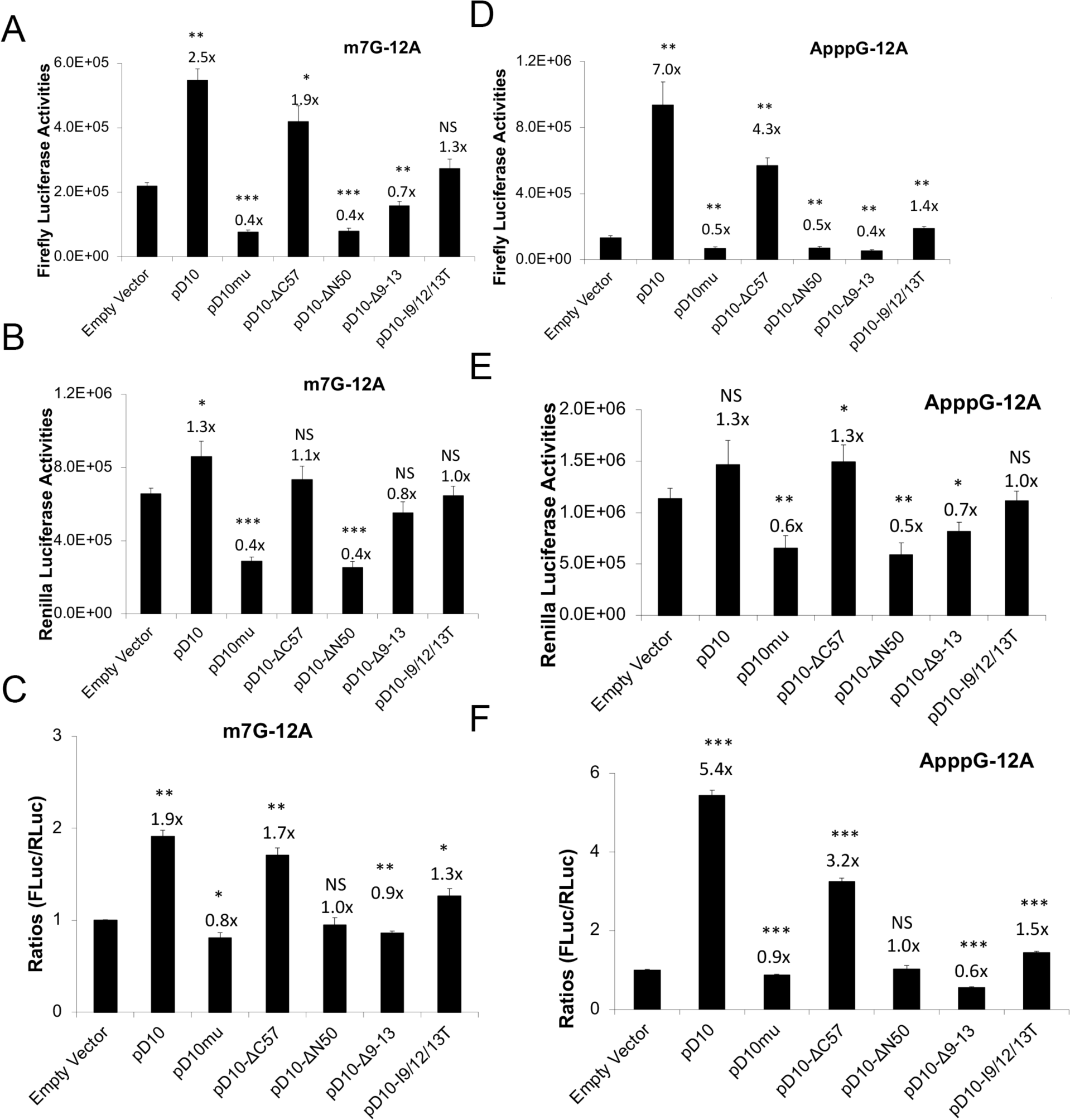
Loss of mitochondrial targeting reduces D10’s mRNA translation enhancement ability for both cap-dependent and cap-independent translation. **(A-C)** 293T cells were transfected with indicated plasmids. 42 h post-transfection, *in vitro* synthesized, m^7^G-capped 12A-Fluc and Kozak-Rluc were co-transfected into the 293T cells. Luciferase activities were measured 6 h post RNA transfection. Fluc **(A),** Rluc **(B)**, and Fluc/Rluc ratios with the empty vector normalized to 1 **(C)** are presented. **(D-F)** 293T cells were transfected with indicated plasmids. 42 h post-transfection, *in vitro* synthesized, ApppG-capped 12A-Fluc and Kozak-Rluc were co-transfected into the 293T cells. Luciferase activities were measured 6 h post RNA transfection. Fluc **(D),** Rluc **(E)**, and Fluc/Rluc ratios with the empty vector normalized to 1 **(F)** are presented. Error bars represent the standard deviation of three replicates. Significance determined by students *t-test* where p>0.05 (ns), p≤0.05 (*), p≤0.01 (**), p≤0.001 (***). The numbers above significance represent fold changes. Significance and fold changes were compared to the empty vector.

## Discussion

In this study, we identified and characterized the first mitochondria-localized decapping enzyme, D10, encoded by a poxvirus, which is required for its unusual function to promote 5’-poly(A)-leader-mediated mRNA translation, including cap-independent translation enhancement. The mitochondrial localization is also necessary for D10’s function to efficiently remove 5’-cap of RNAs and shut off gene expression. Consequently, mitochondrial localization is required for efficient VACV replication. We also pinpointed three hydrophobic Isoleucine residues at the N-terminus of D10 that are essential for D10’s mitochondrial localization. While we do not know exactly if D10 is “in” or “on” the mitochondria, it likely resides on mitochondria such that its catalytically active Nudix decapping motif can be exposed to the cytoplasm to remove 5’-caps of mRNAs. Further study will investigate this aspect.

There are several non-exclusive mechanisms by which mitochondrial localization is required for the optimal effect of D10 to shut off gene expression (**Fig 8**). First, D10 may need to assemble decapping and/or mRNA degradation complex comprising other cellular or viral proteins on mitochondria for function. Second, parking on mitochondria efficiently concentrates D10 locally, which could amplify the decapping efficiency of D10. In fact, mounting evidence shows that many proteins can concentrate for maximal effects, such as phase seperation [45]. For decapping enzymes, one model of Dcp2 function is through concentrating decapping and RNA degradation complex in P-bodies for efficient mRNA decay [14, 15, 46]. Third, mitochondria serve as the highly dynamic vehicles to transport D10 throughout the cytoplasm to more readily access mRNAs, as mitochondria are highly mobile organelles to perform their functions [47, 48]. Fourth, mitochondrial localization may be required for proper conformation of D10 to remove RNA 5’-cap efficiently. Further investigations of these possibilities are ongoing.

**Fig 8.**
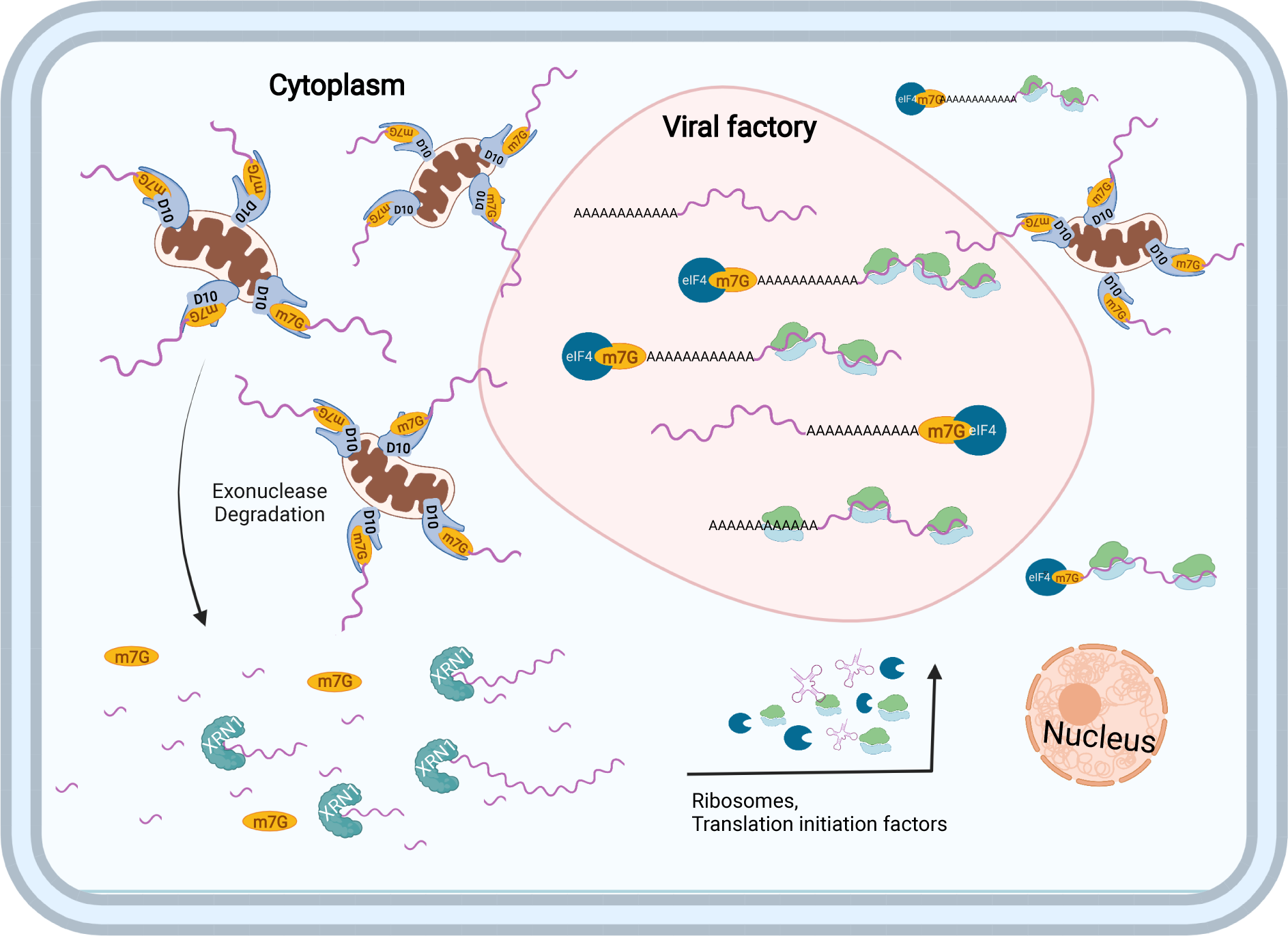
A model on how D10 mitochondria localization impacts its functions. By localizing to mitochondria, D10 (I) preferentially decaps cellular cytoplasmic mRNAs, (II) concentrates locally for powerful decpping activity, (III) rapidly mobilizes in the cytoplasm to access RNA substrates, (IV) assembles decapping and mRNA degradation complex, and (V) frees up and restricts its competition with translation machinery.

How D10’s mitochondrial localization facilitates mRNA translation, especially cap-independent translation enhancement, is thought-provoking but needs extensive investigation of D10’s molecular functions in the presence and absence of VACV infection. As inactivation of its decapping activity also renders it to lose its ability to promote translation [35], the mitochondrial localization requirement for translation promotion could be partially due to its substantially reduced ability to induce mRNA degradation to release translation machinery. In the meantime, the mitochondrial localization restricts its ability to interfere with ribosome recruitment by mRNAs, either in a cap-dependent or a cap-independent manner. Because both decapping activity and mitochondria location are needed for translation promotion, these two functions likely promote mRNA translation in a synergistic manner.

In addition to being required for optimal gene expression shutoff and translation promotion, D10’s mitochondrial localization can provide additional mechanisms to promote VACV replication in VACV-infected cells. Notably, it provides a spatial mechanism for D10 to more readily decap cellular mRNAs (**Fig 8**). Viral factories formed during VACV infection characterized by intensive viral DNA staining are the site for viral DNA transcription, where the viral transcripts can also be translated to produce proteins [49]. As mitochondria are rarely found in the viral factories (**Figs. 1&2**), D10 less readily can access viral mRNAs in the viral factories, especially those post-replicative mRNAs transcribed after viral DNA replication. In contrast, cellular mRNAs and early viral mRNAs transcribed before viral factories formation will more likely be accessed by D10 on highly mobile mitochondria for decapping and subsequent degradation. We and others previously observed pervasive transcription initiation of the VACV genome, especially during the late time of replication [50–53]. These transcripts can be from sense and antisense strands of the viral genome. Many of these transcripts are likely small in size but still get capped. More importantly, many of them can form dsRNA. As these RNAs are less likely loaded by ribosomes and get translated, they may be more likely to escape from viral factories to be accessed by D10 on mitochondria (**Figs. 1,2**).

The almost exclusive localization to mitochondria is a true innovative mechanism for a viral decapping enzyme, which separates it from accessing some viral RNAs to destabilize mRNAs and interfere translation. Meanwhile, it can rapidly find cellular mRNAs. This is unqiue among currently know decapping enzymes. Human Dcp2 is predominantly in the cytoplasm, with many concentrated in P-bodies, particularly under stress [14, 15, 19, 37]. NUDT16 mainly localizes in nuclei, especially in nucleoli [21], suggesting its main function is to regulate nucleolar RNAs. A decapping enzyme from Africa Swine Fever Virus mainly localized to the endoplasmic reticulum colocalizes with RNA cap structure [54]. NUDT12 localizes to a few discrete cytoplasmic granules distinct from P-bodies for Cytoplasmic Surveillance of NAD-Capped RNAs [22]. These studies suggests diverse strategies decapping enzymes used for their functions, demanding the further investigation of these fascinating proteins.

In summary, this study identified a spatial mechanism for a poxvirus-encoded decapping enzyme to regulate mRNA metabolism and translation, resulting in a critical role in viral replication. This study also provides a new direction for decapping enzymes, a group of diverse proteins with important physiological functions.

## Material and Methods

### Cells and Viruses

A549 control cells and A549 DKO cells (kind gifts from Dr. Bernard Moss [34]), Human Foreskin Fibroblasts (HFFs) (a kind gift from Dr. Nicholas Wallace), HeLa cells (ATCC CCL-2), 293T (ATCC-CRL-3216), BHK-21 (C-13), were cultured in Dulbecco’s minimal essential medium (DMEM; Quality Biological). BS-C-1(ATCC CCL-26) cells were cultured in Eagles Minimal Essential Medium (EMEM, Quality Biological). The cell culture media were supplemented with 10% fetal bovine serum (FBS; Peak Serum), 2 mM glutamine (Quality Biological), 100 U/ml of penicillin (Quality Biological), and 100 μg/ml streptomycin (Quality Biological). Cells grow at 37°C with 5% CO_2_.

VACV Western Reserve (WR) strain (ATCC VR-1354) is used in this study. Other recombinant VACVs used in this study were derived from VACV WR strain. vD9muD10mu, vD10mu, vΔD10, vΔD9 were kindly provided by Dr. Bernard Moss and described elsewhere [31, 32].vD10-3xFlag expressing VACV D10 with a 3xFlag tag at the C-terminus was described previously [35]. Recombinant VACVs carrying mutant D10, including vD10-Δ9-13, vD10-I9/12/13T, vΔD9-D10-Δ9to13, vΔD9-D10-I9/12/13T, were generated through homologous recombination using DNA fragments carrying indicated mutations, respectively, followed by three to four rounds of plaque purification of the recombinant viruses.

VACV and its derived recombinant viruses were grown in HeLa or A549DKO cells and purified on 36% sucrose gradient. The viruses (except for vD9muD10mu) were titrated using a plaque assay as described elsewhere [55]. The vD9muD10mu was titrated in A549DKO cells as described elsewhere using anti-VACV antibody immune staining [32, 55].

### Virus infection and plaque assay

Virus infection was performed with DMEM or EMEM containing 2.5% FBS. Virus was sonicated and diluted according to the indicated MOI. Medium containing desired amounts of viruses was added to the cultured cells and incubated at 37°C for 1 h and replaced with fresh medium. For plaque assay, virus-containing-samples were 10-fold serial diluted and added on top of BS-C-1 cells in 12-well plates. After 1 h of incubation at 37°C, the medium was replaced with fresh medium containing 0.5% methylcellulose (Fisher Scientific). Plaques were visualized by staining the infected cells in 12-well plates with 20% ethanol containing 0.1% crystal violet for 5 min.

To compare plaque sizes, the diameters of twenty-five representative plaques of each virus were picked and measured with Image J software.

### Antibodies and Chemicals

Mouse *α*-Flag monoclonal antibody (used for Western blotting analysis) was purchased from Sigma-Aldrich (F3165). Rabbit *α*-Flag polyclonal antibody (used immunostaining for confocal microscopy) was purchased from Thermo Fisher Scientific (PA1-984B). Mouse *α*-Tom20 antibody (sc-17764) and mouse *α*-GAPDH antibody (sc-365062 HRP) were purchased from Santa Cruz Biotechnology. Rabbit *α*-VACV and rabbit *α*-A7 antibodies were kindly provided by Dr. Bernard Moss [56]. Mouse *α*-E3 and mouse *α*-D13 antibodies were kindly provided by Dr. Yan Xiang [57]. Mouse *α*-Cap antibody (201-001) was purchased from Synaptic Systems. MitoTracker (M7510) was purchased from Thermo Fisher Scientific.

### RNA extraction and qRT-PCR

Trizol reagent (Fisher Scientific, 15-596-018) was used for RNA extraction following product instructions. Five μg of RNA was used for reverse transcription with SuperScript III Reverse Transcriptase kit (Fisher Scientific, 18-080-044) following product instructions, using random hexamers as the primers. Quantitative PCR was performed using All-in-one qPCR Mix (GeneCopoeia, QP005) with primers specific for E3L, D13L, A7L, GADPH, and 18s rRNA.

### Plasmids and transfection

Plasmids encode D10 mutants are illustrated in Fig 2A and include: pD10-ΔN8, pD10-ΔN13, pD10-ΔN34, pD10-ΔN50, pD10-ΔC57, pD10-Δ9-13, pD10-I9/12/13T. These plasmids were generated using Q5 Site-Directed Mutagenesis Kit (New England Biolabs, E0554) instructions based on the previously described codon-optimized pD10-3xFlag according to the manufacturer’s [35]. According to the manufacturer’s instructions, plasmid transfection was carried out using lipofectamine 2000 (ThermoFisher scientific, 11668019).

### Western Blotting Analysis

Western blot was performed as described previously [58]. Briefly, the samples were resolved by sodium dodecyl sulfate-polyacrylamide gel electrophoresis (SDS–PAGE), followed by transferring to a polyvinylidene difluoride membrane (PVDF). The PVDF membrane was blocked with 2% BSA or 5% milk plus 1% BSA at room temperature for 1 h, then incubated with primary antibodies in the same blocking buffer at room temperature for 1 h or at 4°C overnight. After washing with TBST three times, membranes were incubated with secondary antibodies at room temperature for 1 h and washed three times with TBST. Before imaging, the membrane was developed using SuperSignal West Femto Maximum Sensitivity Substrate (Thermo Fisher Scientific, 34094). Antibodies were stripped from the membrane by Restore Western Blot stripping buffer (Thermo Fisher Scientific, 21059) for analysis using another antibody.

### Immunostaining and confocal microscope

Mock, VACV-infected or-plasmids transfected cells were fixed with 4% paraformaldehyde solution for 30 min at room temperature. The cell membrane was penetrated with 1xPBS containing 0.5% TritonX-100 for 10 min following being blocked with 1xPBS containing 2% BSA for 1 h. Primary antibodies were diluted in PBS (with 2% BSA) and incubated with cells for 1 h at room temperature. After three times of washing with 1xPBS, cells were incubated with secondary Alexa Fluor (488 nm for green and 594 nm for red)-conjugated IgG diluted in 1x PBS (with 2% BSA) at room temperature for 1 h. After three times of washing with 1xPBS, cells were stained with DAPI for 5 min and washed with 1xPBS two more times. Coverslips were mounted using 40% glycerol. Zeiss 880 or Zeiss 700 confocal microscopy was used to visualize the cells.

### *In vitro* RNA synthesis, transfection, and luciferase assay

Synthesis of RNA *in vitro* was carried out as previously described using HiScribe T7 Quick High Yield RNA Synthesis Kit (New England Biolabs, E2050) [35, 41, 42]. The RNAs were co-transcriptionally capped with m^7^G anti-reverse cap analog or ApppG Cap Analog (New England Biolabs, 1411 and Cat#1406). The RNAs were purified using a Purelink RNA Mini Kit (Thermo Fisher Scientific, 12183025) and transfected into cells using Lipofectamine 2000 (Thermo Fisher Scientific, L11668019) according to the manufacturer’s instructions. Six hours post-transfection, cell lysates were collected, and luciferase activities were measured using a Dual-Luciferase Reporter Assay System (Promega, E1960) and GloMax Navigator Microplate Luminometer with dual injectors (Promega) as per manufacturer protocol.

### *Gaussia* luciferase assay

The *Gaussia* luciferase activities were measured using a luminometer using the Pierce Gaussia luciferase flash assay kit (Thermo Scientific, 16158).

### Statistical analysis

The student’s *t-*test was performed to evaluate statistical differences from at least three replicates. We used the following convention for symbols to indicate statistical significance: ns, *P* > 0.05; *, *P* ≤ 0.05; **, *P* ≤ 0.01; ***, *P* ≤ 0.001; ****, *P* ≤ 0.0001.

## Acknowledgment

We thank Bernard Moss, Yan Xiang, and Nicholas Wallace for providing reagents and materials. We thank Joel Sanneman at Kansas State University CVM Confocal Facility for technical assistance. ZY is supported by a grant from the National Institutes of Health (R01 AI143709).

---

**Fig S1.**
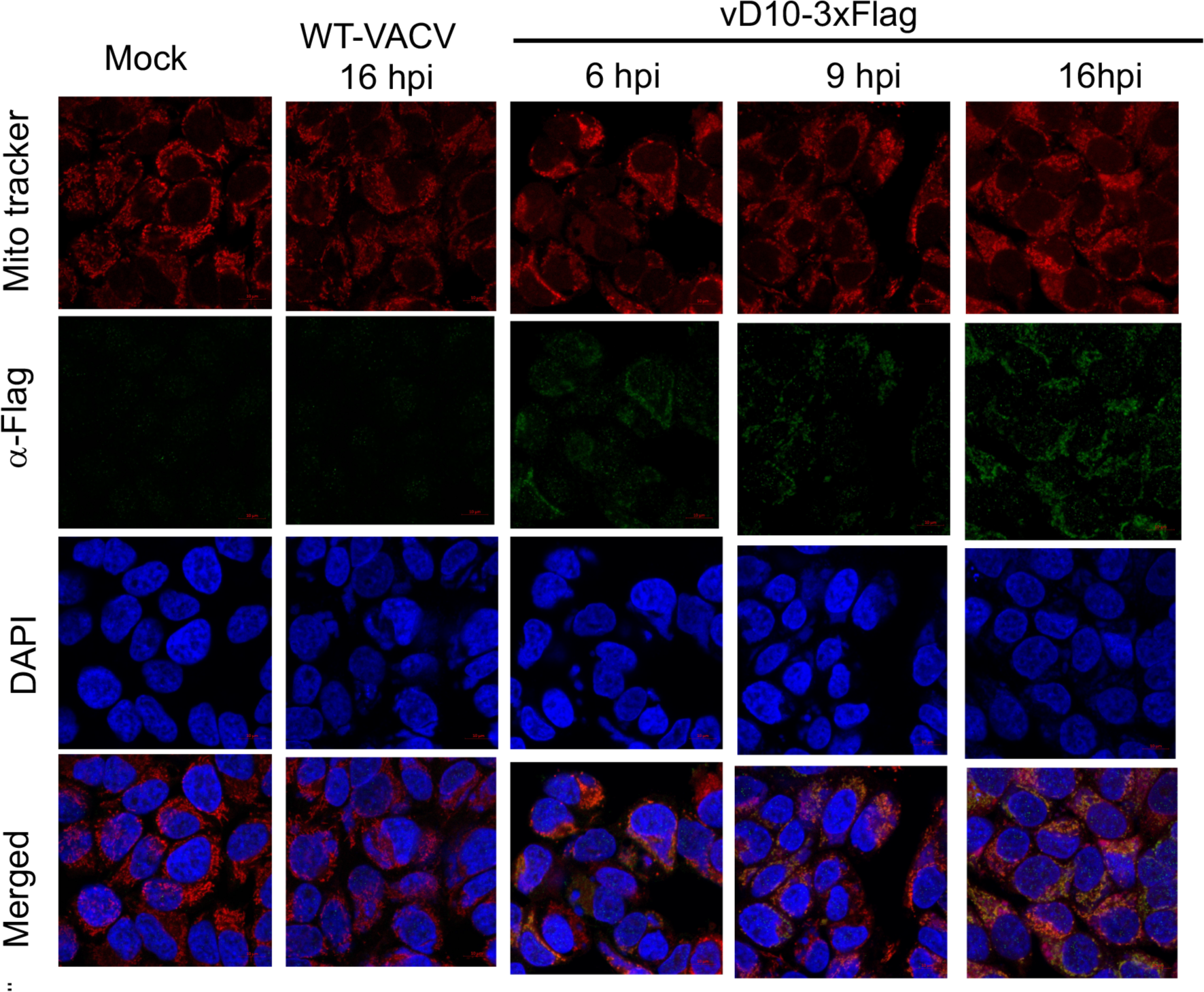
D10 localizes to mitochondria in HeLa cells during infection. HeLa cells were infected with vD10-3xFlag, or WT-VACV, or mock-infected. Confocal microscopy was used to visualize D10 (*α*-Flag antibody, green), mitochondria (MitoTracker, red), and DNA (DAPI, blue) at 6, 9, and 16 hpi.

**Fig S2.**
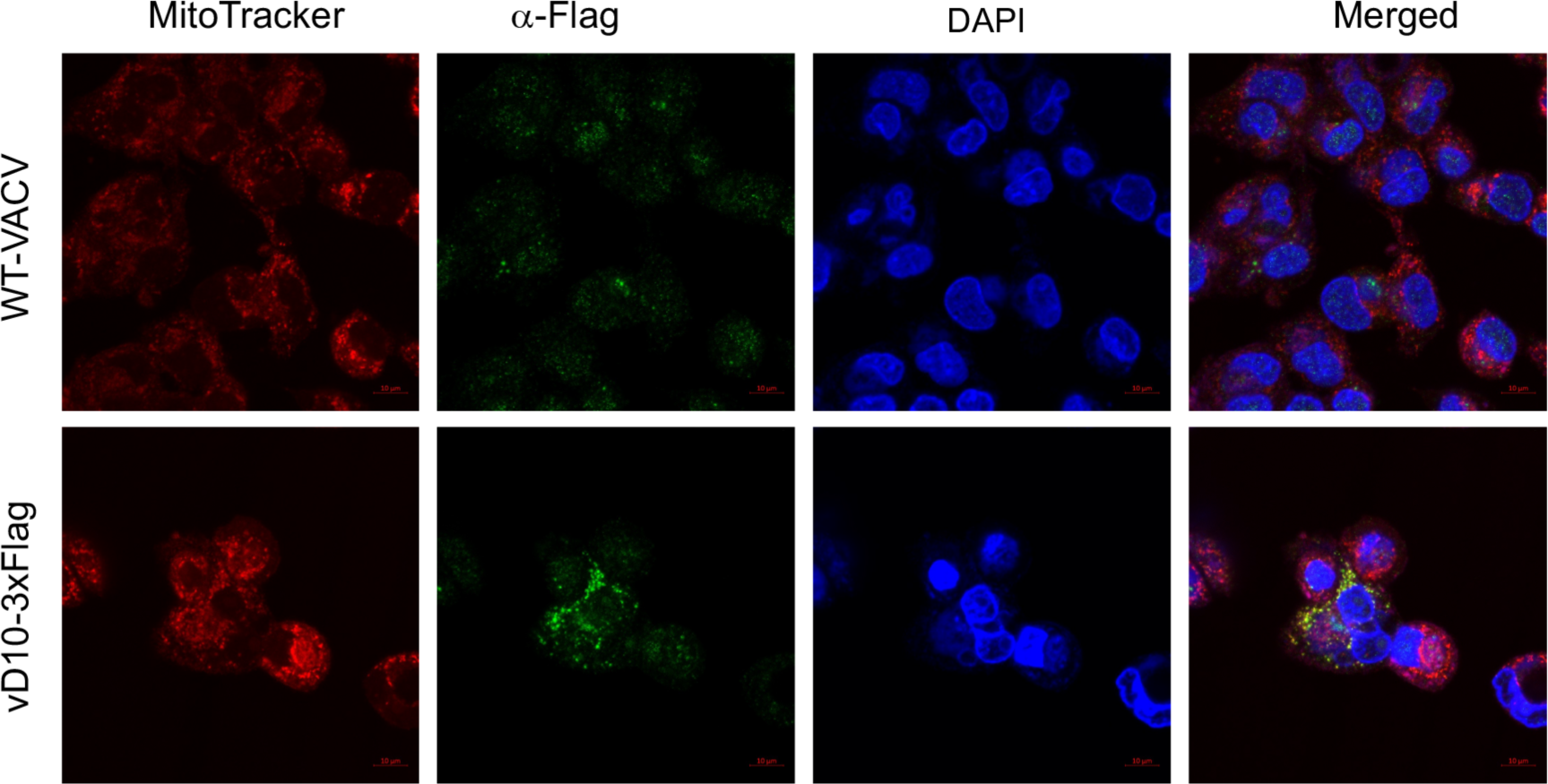
D10 localizes to mitochondria in A549DKO cells during infection. A549DKO cells were infected with vD10-3xFlag or WT-VACV. Confocal microscopy was employed to visualize D10 (*α−*Flag antibody, green), mitochondria (MitoTracker, red), and DNA (DAPI, blue) at 16 hpi.

**Fig S3.**
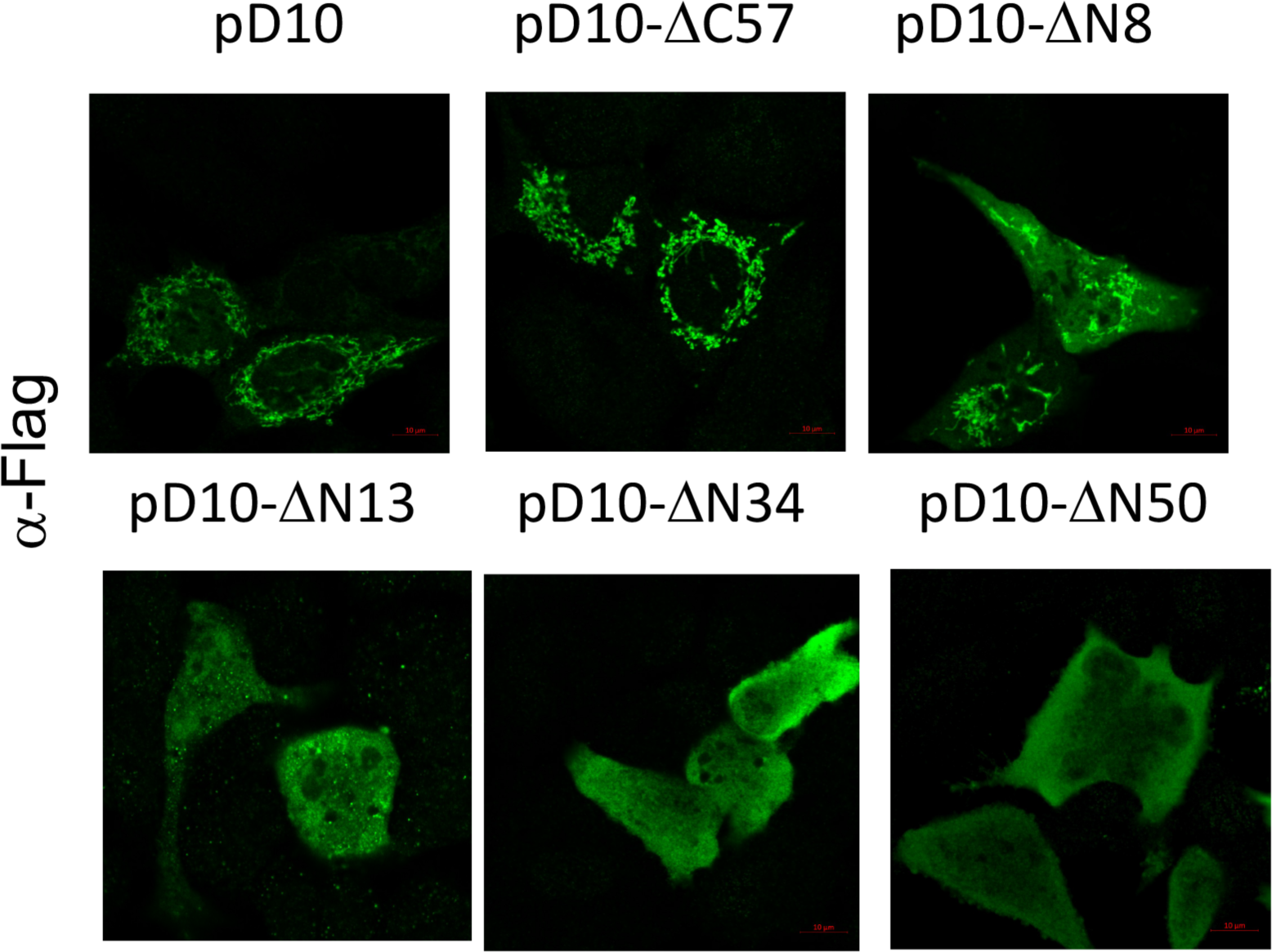
D10 mitochondrial signal is located at the N-terminus of D10. A549 DKO cells were transfected with a plasmid expressing indicated codon-optimized D10 truncation mutants with C-terminal 3xFlag. Confocal microscopy was employed to visualize D10 or its mutants using *α−*Flag antibody (green).

**Fig S4.**
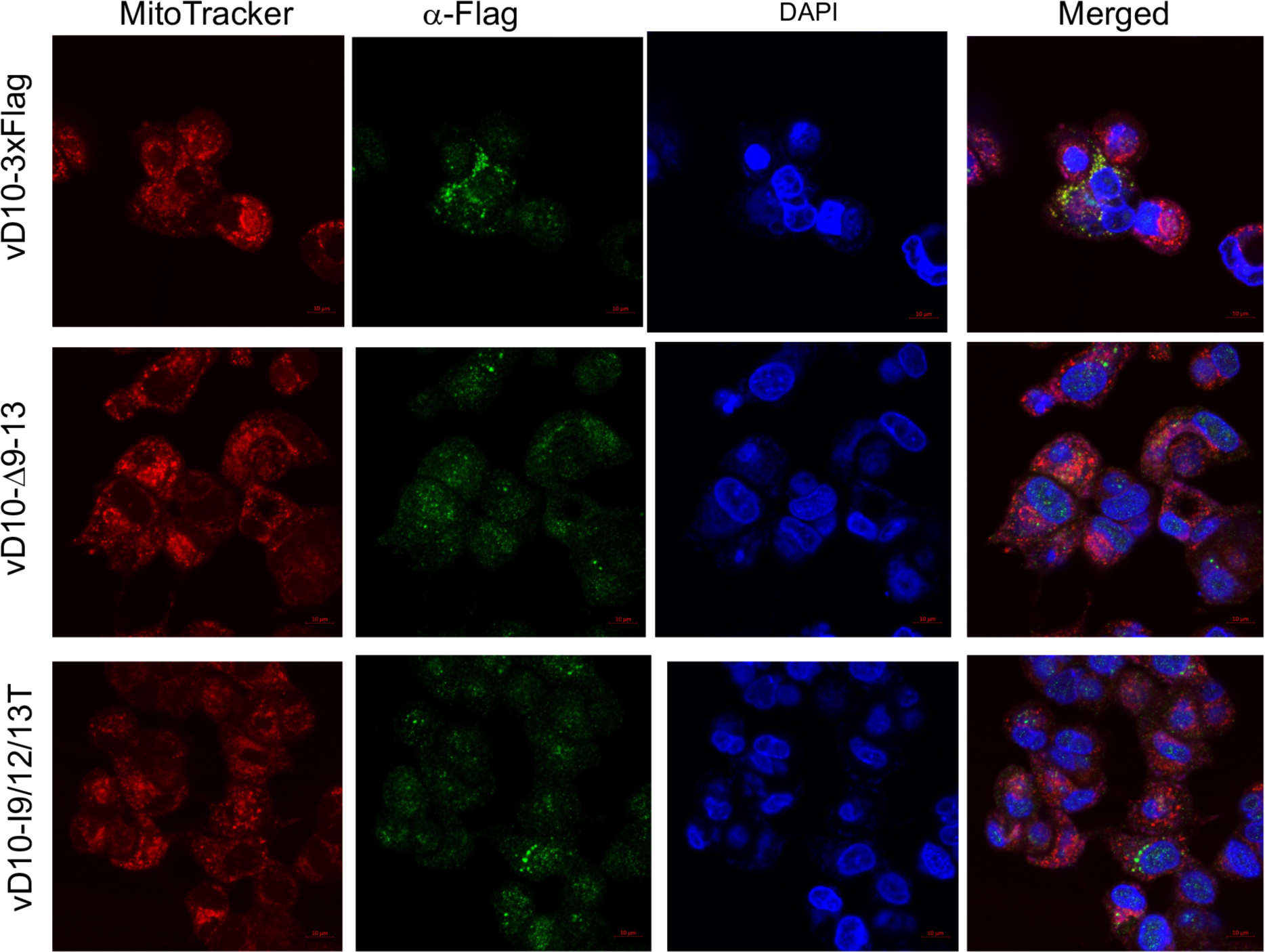
D10 mutants with amino acids 9-13 deletion or mutation expressed from recombinant VACV do not localize to mitochondria during infection, A549DKO cells were infected with indicated recombinant VACVs (MOI=3) encoding D10 mutants with a C-terminal 3xFlag tag. Confocal microscopy was used to visualize D10 (*α*-Flag antibody, green), mitochondria (MitoTracker, red), and DNA (DABI, blue) at 16 hpi.

**Fig S5.**
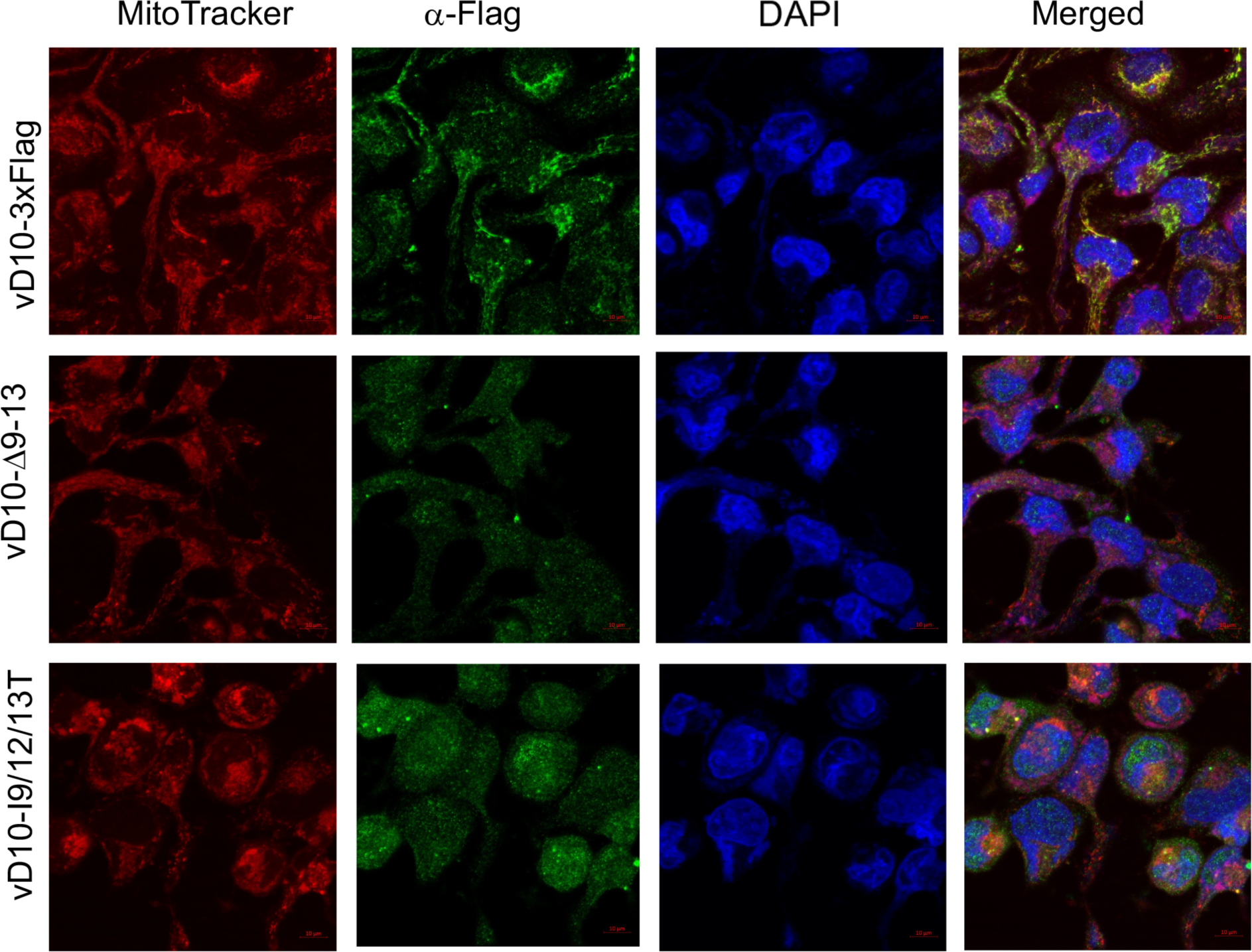
D10 mutants with amino acids 9-13 deletion or mutation expressed from recombinant VACV do not localize to mitochondria during infection. HeLa cells were infected with indicated recombinant VACVs (MOI=3) encoding D10 mutants with a C-terminal 3xFlag tag. Confocal microscopy was used to visualize D10 (*α*-Flag antibody, green), mitochondria MitoTracker, red), and DNA (DAPI, blue) at 16 hpi.

**Fig S6.**
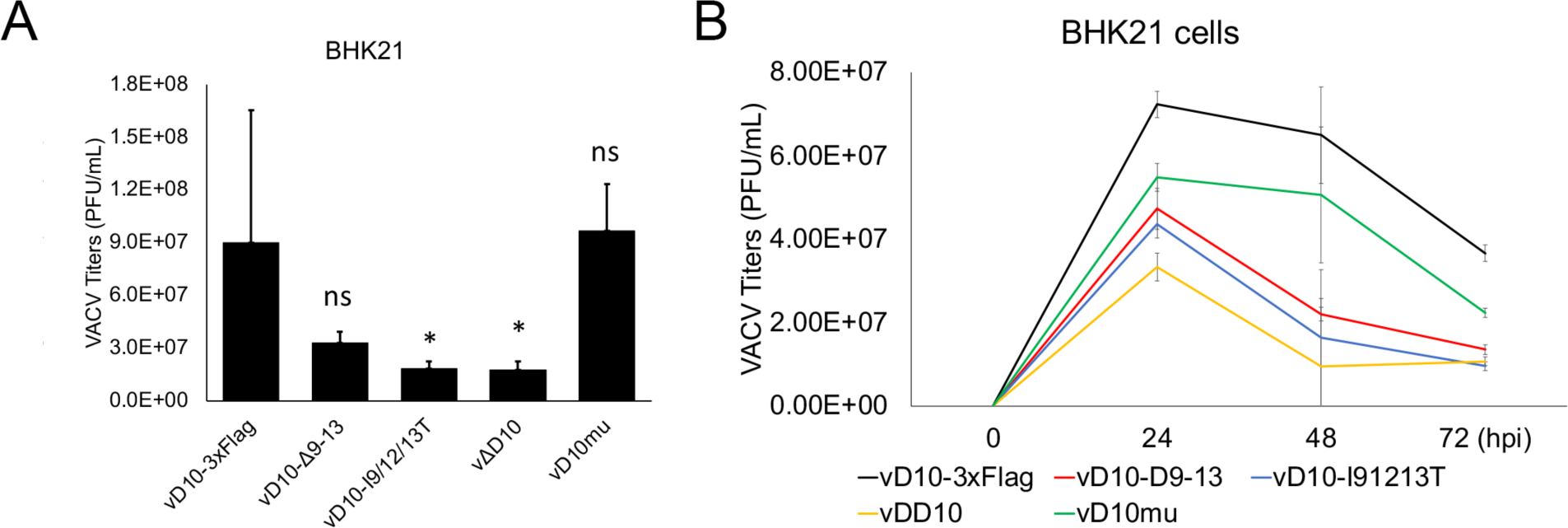
BHK-21 cells were infected with indicated viruses at an MOI of 3 (**A**) or 0.001 (**B**). Viral titers were determined using a plaque assay at indicated times post-infection. All the viruses used encode D9. Error bars represent the standard deviation of at least three replicates. ns, P > 0.05; *, P ≤ 0.05. Significance was compared to vD10-3xFlag.

